# Immunogenicity and tolerability of a SARS-CoV-2 TNX-1800, a live recombinant poxvirus vaccine candidate, in Syrian Hamsters and New Zealand White Rabbits

**DOI:** 10.1101/2023.09.06.556620

**Authors:** Mayanka Awasthi, Anthony Macaluso, Scott J Goebel, Erin Luea, Ryan S Noyce, Farooq Nasar, Bruce Daugherty, Sina Bavari, Seth Lederman

**Affiliations:** Tonix Pharmaceutical, Frederick, MD, USA; Southern Research, Birmingham, AL, USA; University of Alberta, Department of Medical Microbiology & Immunology and Li Ka Shing Institute of Virology, University of Alberta, Edmonton, Alberta, Canada; Tonix Pharmaceuticals, Chatham, NJ; Tonix Pharmaceuticals, Dartmouth, MA, USA

**Keywords:** SARS-CoV-2, TNX-1800, vaccine, vaccine platform, horsepox, spike protein, antibody titer, immunogenicity

## Abstract

TNX-1800 is a preclinical stage synthetic derived live chimeric horsepox virus vaccine that comprises an engineered SARS-CoV-2 spike (S) gene expression cassette. The objectives of this study were to assess the immunogenicity and tolerability of TNX-1800 administration in Syrian golden hamsters and New Zealand white rabbits. Animals were vaccinated via percutaneous inoculation and evaluated for dose tolerance and immunogenicity at three different dose levels. The 28-day study data showed that the single percutaneous administration of three TNX-1800 vaccine dose levels was well tolerated in both hamsters and rabbits. For all dose levels, rabbits had more dermal observations than hamsters at the same dose levels. Vaccine-induced viral load four weeks post-dosing was below the detection level for both species.

## 1. Introduction

The COVID-19 pandemic, caused by the novel coronavirus SARS-CoV-2, has had a profound impact on global public health and the world economy. The virus first emerged in Wuhan, China, in late 2019 and has since spread rapidly, leading to more than 10 million deaths and widespread social disruption [1-3]. In response, scientists and pharmaceutical companies worldwide have been working tirelessly to develop safe and effective vaccines to prevent COVID-19 infection and reduce its transmission. The development of COVID-19 vaccines has been a massive undertaking, involving unprecedented levels of collaboration and innovation. Several different types of vaccines have been developed, including mRNA vaccines [4], live vaccines [5], inactivated vaccines [6], viral vector vaccines [7], and protein subunit vaccines [8]. These vaccines have undergone rigorous clinical trials to ensure their safety and efficacy with some of these vaccines receiving US Food and Drug Administration (FDA) approval. Despite the remarkable success of COVID-19 vaccine development, the global rollout of vaccines has faced significant challenges. Limited vaccine supply, distribution logistics, and vaccine hesitancy have all contributed to disparities in vaccine access and uptake [9]. As the SARS-CoV-2 infection continues to evolve and with the emergence of new variants, the development, and distribution of new vaccines remain critical to reduce the impact of SARS-CoV-2 infection. Specifically, the development of vaccines that do not require ultra-cold storage and can be given one time with the ability to protect against multiple variants would make distribution much easier and cheaper in resource-limited settings. Continued research and innovation are needed to address the ongoing challenges of vaccine access and hesitancy, as well as to develop new vaccines that can provide protection against emerging future variants of the virus.

TNX-1800 is a preclinical stage synthetic derived live chimeric horsepox virus vaccine that comprises an engineered SARS-CoV-2 S gene expression cassette. The horsepox virus (HPXV), being a phylogenetically close relative of the modern vaccinia virus (VACV) and other viruses from which the core vaccines are generated, provides a possible alternative to produce vaccines for other infectious diseases [10]. Developed as a live horsepox virus, TNX-1800 has been modified to express the SARS-CoV-2 spike protein for stimulation of an immune response against SARS-CoV-2. One of the initial steps in developing TNX-1800 is a better understanding of its tolerability and immunogenicity in small animal models. In this report, we have assessed TNX-1800 immunogenicity, tolerability, and vaccine systemic dissemination in Syrian golden hamsters and New Zealand white rabbits, when administered by percutaneous administration.

Animals were monitored based on clinical observations, local Draize scoring, body weights, antibody production, and viral load in skin samples collected from the injection sites at necropsy. Single percutaneous administration of various TNX-1800 vaccine dose levels was well tolerated in hamsters and rabbits. SARS-CoV-2 spike protein-specific Immunoglobulin G (IgG) titers indicated a higher response in rabbits receiving TNX-1800 than hamsters receiving the same vaccine dose. All TNX-1800 vaccinated hamsters and rabbits maintained circulating SARS -CoV-2-specific antibodies until the study end, indicating a minimum weeklong-four-week-long vaccination induced immune response. Vaccine-induced viral load assessed by qPCR in skin samples collected at the dose administration site four weeks postdosing was below the detection level for both animal species in all groups. The encouraging data on tolerability and immunogenicity of TNX-1800 in small animal models have paved the path to continue testing the vaccine in nonhuman primates. The nonhuman primate studies will include tolerability, safety, and a deeper understanding of cellular and humoral responses to protective antigens of SARS-CoV-2 and other infectious diseases.

## 2. Materials and Methods

### 2.1. Vaccine Information

TNX-1800 is a recombinant HPXV [10], expressing the codon-optimized, SARS-CoV-2 Spike gene (Wuhan strain, NC_045512) from a non-essential insertion locus. Expression of the SARS-CoV-2 Spike gene is driven by a synthetic early/late poxvirus promoter. The methods for the generation of the recombinant VACV control vaccine, TNX-1200, have been previously described (PMID 35290406). Both virus preparations were resuspended in Tris-HCl (10 mM, pH 8.0).

### 2.2. Ethics Statement

This work was supported by an approved Institute Animal Care and Use Committee (IACUC) animal research protocol in compliance with the Animal Welfare Act, PHS Policy, and other Federal statutes and regulations relating to animals and experiments involving animals. The facility where this research was conducted is accredited by the Association for Assessment and Accreditation of Laboratory Animal Care (AAALAC International) and adheres to principles stated in the Guide for the Care and Use of Laboratory Animals, National Research Council, 2011[11].

### 2.3 Study Design and Immunization Procedure

Syrian golden hamsters ages at least 13 weeks and weighing ∼80-150 g and New Zealand white rabbits ages at least 13 weeks and weighing 2.0 to 3.5 kg were obtained. The hamsters and rabbits were placed into four individual study groups (groups 1-4) and the rabbits (groups 1-4) as shown in **Figure 1A**. All animals were immunized on day 1 via percutaneous inoculation with identified doses for each group (**Figure 1A**). Briefly, the vaccination site was located between the shoulder blades. Prior to each dose administration, the dosing site was shaved and marked. The site was sanitized, and 10 μL of either TNX-1200 or TNX-1800 was placed on the skin using a sterile pipette tip. A bifurcated needle was used to push each virus vaccine under the skin by penetrating the skin vertically approximately 15 times. The residual vaccine was removed using sterile gauze.

**Figure 1.**
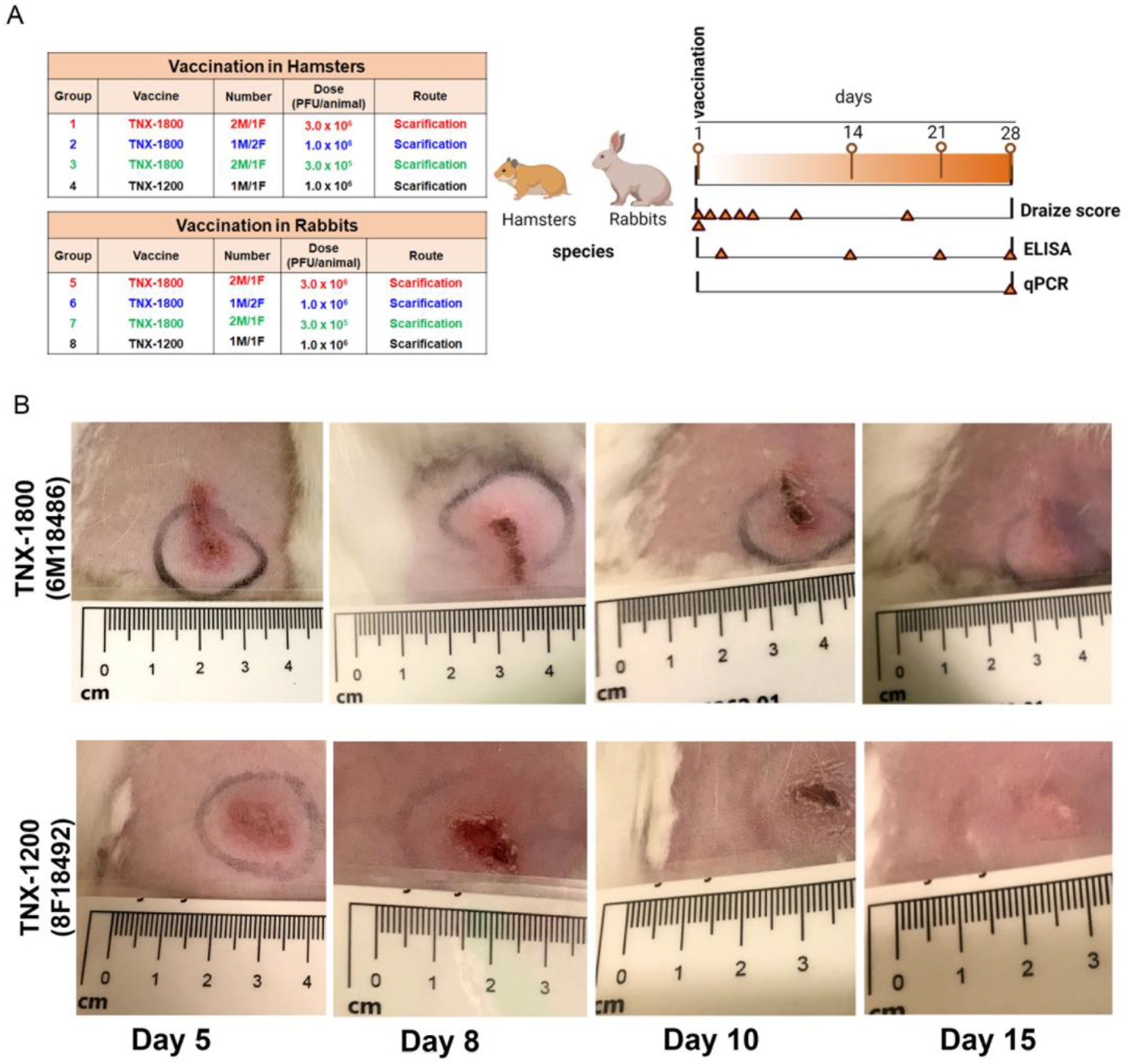
**(A)** Schematic of the hamsters and rabbits study design. Males (M) and females (F) hamsters and rabbits were vaccinated with TNX-1800 (three different dose levels), TNX-1200. Groups 4 and group 8 were used as a positive control for erythema and edema scoring. Following vaccination, Draize score, immunogenicity and qPCR was performed at intervals indicated by triangles. **(B)** Photographs of the lesions at the injection site of Group 6 (TNX-1800) and Group 8 rabbits (TNX-1200).

### 2.4. Body Weight, Lesion Counts, and Draize Scoring

Body weight and detailed clinical observations specifically including evaluations of erythema, edema, and percutaneous (eschar/scab/pox formation) were determined starting, prior to dosing on Day 1, and twice weekly. Photos were taken of all lesions and any abnormal skin observations.

### 2.5. Enzyme-Linked Immunosorbent Assay (ELISA)

Serum samples were collected during the vaccination phase as per the time point shown in **Figure 1** and were used to determine binding antibody titers specific for SARS-CoV-2 spike protein. Serum samples were collected on days 1, 14, 21, and 28. To measure SARS-CoV-2 spike protein antibody responses during the vaccination phase, serum samples were analyzed using a validated SARS-CoV-2 spike protein-specific ELISA in accordance with Southern Research standard operating procedures. Briefly, 96-well ELISA plates were coated with 100 uL/well of 0.5 ug/mL purified SARS-CoV-2 (2019-nCoV) Spike Protein (Wuhan variant S1+S2 ECD) antigen in phosphate-buffered saline (PBS) and incubated overnight at 2 - 8 °C. Plates were washed using an automated plate washer (BioTek ELx405, Winooski, VT) and then blocked with 5% non-fat milk in 0.05% PBS-Tween 20 (PBST) at 37 ± 1 °C. Serum samples were serially diluted in 5% non-fat milk in 0.05% PBST and added at 100 uL/well. Specific antibodies were detected using a goat anti-Hamster IgG (H+L) Cross-Adsorbed Secondary Antibody, HRP (Invitrogen, Cat# PA5-33286) and Goat anti-Rabbit IgG (H+L) Cross-Adsorbed Secondary Antibody, HRP (Invitrogen, Cat# 31462). Plates were developed with 100 uL/well of ABTS ((2,2’-Azinobis [3-ethylbenzothiazoline-6-sulfonic acid]-diammonium salt) substrate for 15 to 20 minutes, stopped with 100 uL of 1% sodium dodecyl sulfate (SDS) in distilled water and read at 450 nm using a Spectra-Max plate reader (Molecular Devices. Sunnyvale, CA).

### 2.6. Viral Load by Quantitative PCR (qPCR) Analysis

Viral loads were measured in tissue homogenates of skin punch biopsy samples. Total DNA was extracted from each sample using QIAcube robot and viral genome copies were quantified via qPCR. Briefly, samples were extracted using TriZol reagent (Sigma Aldrich) following the manufacturer’s recommendations. Viral DNA isolated from samples was eluted with nuclease-free H_2_O and stored at -70°C or below. The quantitative PCR (qPCR) reaction used to assess vaccine viral load, was targeted to the Orthopox E9L DNA Polymerase gene using commercially available Pan-Orthopox Virus E9L Gene-Specific Quantitative PCR Assay Detection Kit (BEI Resources Cat# NR-9350).

### 2.7 Statistical Analysis

Comparative statistical analysis of the body weight data and other parameters as deemed appropriate were performed using Provantis®. Statistical analysis of other data (e.g., Immunology) consisted of descriptive statistics such as mean, standard deviation, and coefficient of variation.

## 3. Results

### 3.1. Post-Dosing Clinical Observations

To evaluate the immunogenicity and tolerability of TNX-1800, three groups (group 1-3) of hamsters (N=3) and rabbits (n=3) were vaccinated at day 1 via percutaneous administration using a bifurcated needle with different doses of TNX-1800 (**Figure 2A**). Following percutaneous dose administration, animals in groups 1-8 were observed for local dermal responses following vaccination. Photographs were taken of lesions at the injection site: on day 5 and 6 (a male rabbit from group 6 and both group 8 rabbits - a male and a female) and on day 10 and 15 (all rabbits from groups 5-8), representative photographs are shown in **Figure 1B**. Group 4 hamsters and group 8 rabbits, vaccinated via percutaneous inoculation with a single dose of 1.0 x 10^6^ PFU of TNX-1200, were used as a comparative control for erythema and edema scoring. All animals in the study consistently maintained or gained body weights and any fluctuations were considered normal for all the animals (**Table S1**). There were no abnormal clinical observations documented at any dose level for rabbits or hamsters. All hamsters and rabbits exhibited minimal to moderate erythema within 2 hours of dosing, while three rabbits and none of the hamsters had edema. Minimal erythema was resolved in five hamsters and one rabbit 5 hours post-vaccination. Observations of minimal edema increased in frequency in both species beginning on day 2 and continuing through day 5 for hamsters and day 10 for rabbits. The rabbits at all dose levels consistently had more observations related to erythema and edema than hamsters at the same dose levels (**Table S2**).

**Figure 2.**
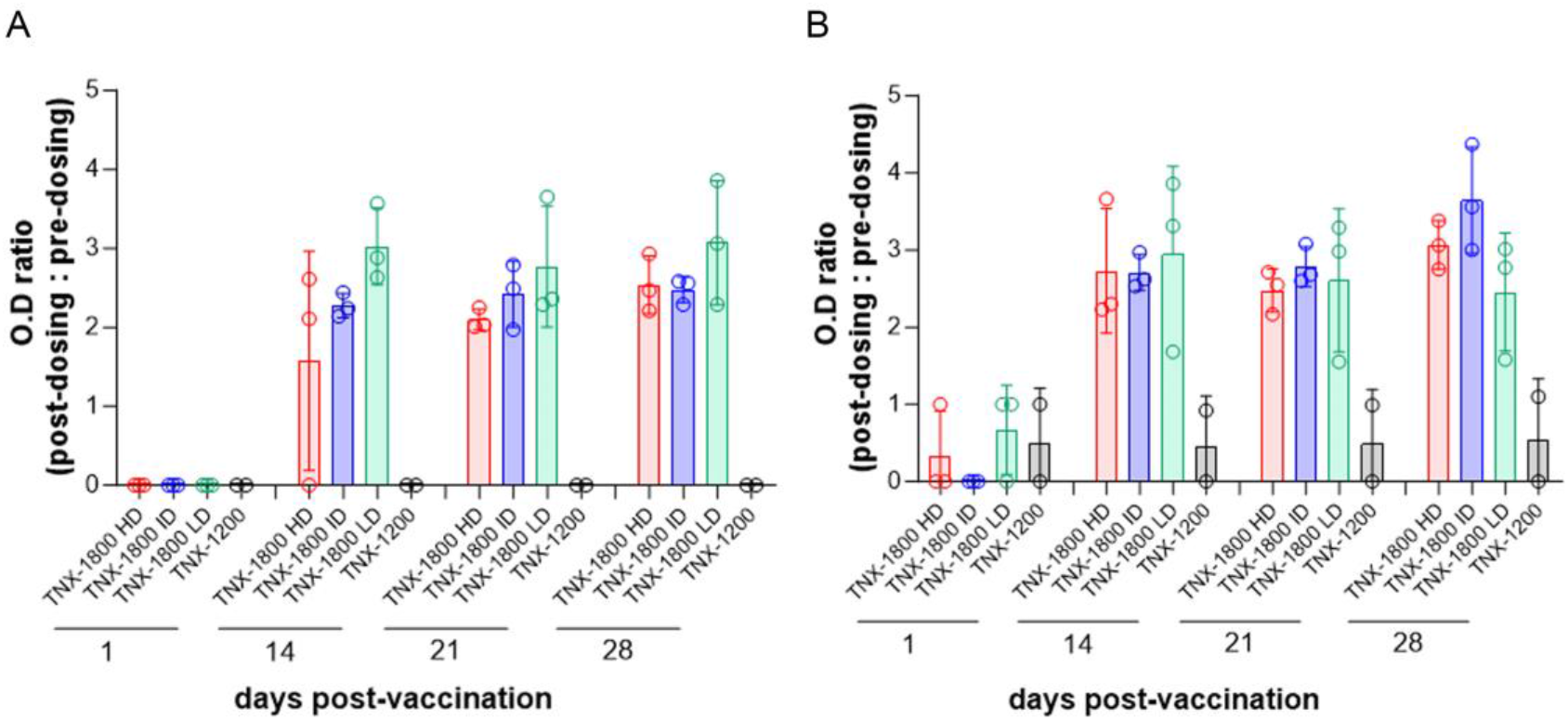
Immunogenicity following vaccination in **(A)** hamsters and **(B)** rabbits as determined by ELISA. Graph depicts the post-dosing to pre-dosing OD ratio corresponding to the total spike specific IgG. HD: High dose (3x10^6^), ID: Intermediate dose (1x10^6^), LD: Low dose (3x10^5^).

### 3.2. TNX-1800 Immunogenicity

Humoral responses in hamster (groups 1-4) and rabbit (groups 5-8) serums were measured via detecting the binding of SARS-CoV-2 spike protein-specific immunoglobulins G (IgG) in an ELISA format. For both species, the titers were relatively unaffected by dose, with all groups quickly reaching the maximal reading by day 14 to 21 post vaccination (**Figure 2A and 2B**).

For high and intermediate dose groups, IgG titer and OD titer ratio values indicated a higher response in rabbits than hamsters. The nine vaccinated hamsters from groups 1-3 generated spike-specific IgGs by day 14 and all TNX-1800 vaccinated hamsters showed antibody production on day 21. IgG titers detected in group 1 peaked on day 21, while the highest values of IgG titers were detected in groups 2 and 3 on day 28. However, these levels of anti-SARS-CoV-2 spike IgG were not statistically different than what was measured at low dose of the vaccine. As expected, control groups vaccinated with TNX-1200 lacking the SARS-CoV-2 spike gene cassette (groups 4 and 8), had low background levels of anti-SARS-CoV-2 spike-specific IgGs throughout the study. Hamsters in group 3 (TNX-1800 dose of 3.0 x 10^5^) consistently had higher average OD titer ratios throughout the study (**Figure 2A**). TNX-1800 vaccinated rabbits consistently had a higher IgG titer value beginning on day 14 when compared to hamsters at the same dose level. Groups 5 and 6 showed the highest average OD titer ratio on day 28: 3.06 and 3.64, respectively. Rabbits in group 7 (TNX-1800 3.0 x 10^5^) had the highest IgG titer ratio on day 14 (2.95) followed by a slight decrease to 2.61 and 2.12 in the following two weeks (**Figure 2B**).

To examine the longevity of TNX-1800 replication at the vaccination site, we tested the site of the vaccine 4 weeks after TNX-1800 administration for the presence of pox-related genes by qPCR. The assessment of the vaccine-induced viral load by qPCR assay in skin samples collected from the injection sites four weeks post-dosing indicated no detectable levels of virus present in the tissues (**Table S2**).

## 4. Discussion

Multiple approaches have been used to deliver SARS-CoV-2 spike as a vaccine and elicit neutralizing immune responses including live viruses, inactivated viruses, viral vectors (eg, adenovirus), nucleic acids (primarily mRNA), and protein subunits [4-8]. The combination of quickly waning immunity, especially in the upper respiratory system, and the introduction of new SARS-CoV-2 variants with modified spike, has raised concern about the current vaccines [12, 13]. Also, accumulating data suggest the currently approved vaccines are not capable to fully protect against the infection because they do not produce ample amounts of neutralizing immunity in the nasal cavity [14]. Thus, there is gap in defense against evolving SARS-CoV-2 infection which may be filled by the ongoing need to optimize SARS-CoV-2 vaccines, either by adapting current vaccine strategies or by searching for alternative technologies. Exploring alternative delivery of SARS-CoV-2 spike as a vaccine such as using novel live vectors may add an additional layer of protection against infection.

Here, we tested the tolerability of a single percutaneous administration of live horsepox vector expressing Wuhan spike gene and its ability to elicit binding antibodies. Various TNX-1800 vaccine dose levels were well tolerated in hamsters and rabbits. Observations of erythema were recorded beginning at 2 hours after administration and edema increased in frequency beginning on day 2. Most local draize observations were noted through day 10. Rabbits at all dose levels consistently had more dermal observations related to erythema and edema than hamsters at the same dose levels. ELISA IgG titer and OD ratio values indicated a higher response throughout the study in rabbits receiving TNX-1800 than in hamsters receiving the same dose levels. Throughout the study, hamsters in Group 3 (TNX-1800, 3.0 x 10^5^ PFU/animal) consistently had higher average OD titer ratios. Although this was not statistically significant. Rabbits consistently had higher IgG titers beginning on day 14 when compared to hamsters at the same dose level. Group average titer ratios detected in groups 5 and 6 peaked on day 28 (3.06 and 3.64, respectively), while the highest group average OD titer ratio was detected in group 7 (TNX-1800 3.0 x 10^5^ PFU/animal) on day 14 (2.95). Post-vaccination viral load was assessed by qPCR in skin samples collected at the dose administration site at four weeks and were found to be below the detection level for both species in all groups (Table A2).

In conclusion, we have performed an in vivo immunogenicity and tolerability study using Syrian golden hamsters and New Zealand white rabbits and demonstrated that the TNX-1800 vaccine is immunogenic in both species. For all dose levels, the average IgG titer values were statistically similar and significantly higher than the control groups (TNX-1200). Future efforts will include testing immunogenicity with lower vaccine doses to minimize the vaccination site reactions as well as the assessment of neutralizing antibody titers against HPXV. These encouraging preliminary immunogenicity and tolerability data suggest the potential of this platform for further development of future vaccines against SARS-CoV-2 and other pandemic-causing viruses.

Future efforts will include testing immunogenicity and efficacy in non-human primates. Additional experiments will allow a better understanding of neutralizing antibody titers and test the ability of the vector to carry multiple protective antigens inserts against various infectious diseases.

## Author Contributions

Conceptualization, A.M., E.L., B.D., and S.B.; methodology, E.L., R.N, ; formal analysis, M.A., A.M., F.N, S.B, S.L.; investigation, M.A., A.M., F.N, S.B, S.L.; resources, A.M., F.N, S.B, S.L.; data curation, M.A., A.M., E.L., F.N, S.B; writing—original draft preparation, M.A., S.B., ; writing— review and editing, M.A., A.M., E.L., R.N., B.D., F.N, S.B, S.L.; visualization, M.A., A.M., F.N, S.B.; supervision, A.M., F.N., S.B., S.L.; project administration, A.M., S.B., S.L.; funding acquisition, S.L. All authors have read and agreed to the published version of the manuscript.

## Institutional Review Board Statement

This study was designed to use the fewest number of animals possible, consistent with the objective of the study, the scientific needs, contemporary scientific standards, and in consideration of applicable regulatory requirements. The study design was reviewed by the Institutional Animal Care and Use Committee (IACUC) at Southern Research and the study was in full compliance with applicable federal and Southern Research requirements. The Institutional Biosafety Committee (IBC) has approved this project. The facility where this research was conducted is accredited by the Association for Assessment and Accreditation of Laboratory Animal Care (AAALAC International) and adheres to principles stated in the Guide for the Care and Use of Laboratory Animals, National Research Council, 2011[11].

## Informed Consent Statement

Not applicable

## Data Availability Statement

Not applicable

## Acknowledgments

The authors wish to acknowledge Southern Research and members of Tonix Pharmaceuticals for providing insightful comments during the conduct of these studies.

## Supplementary Information

**Table S1.**
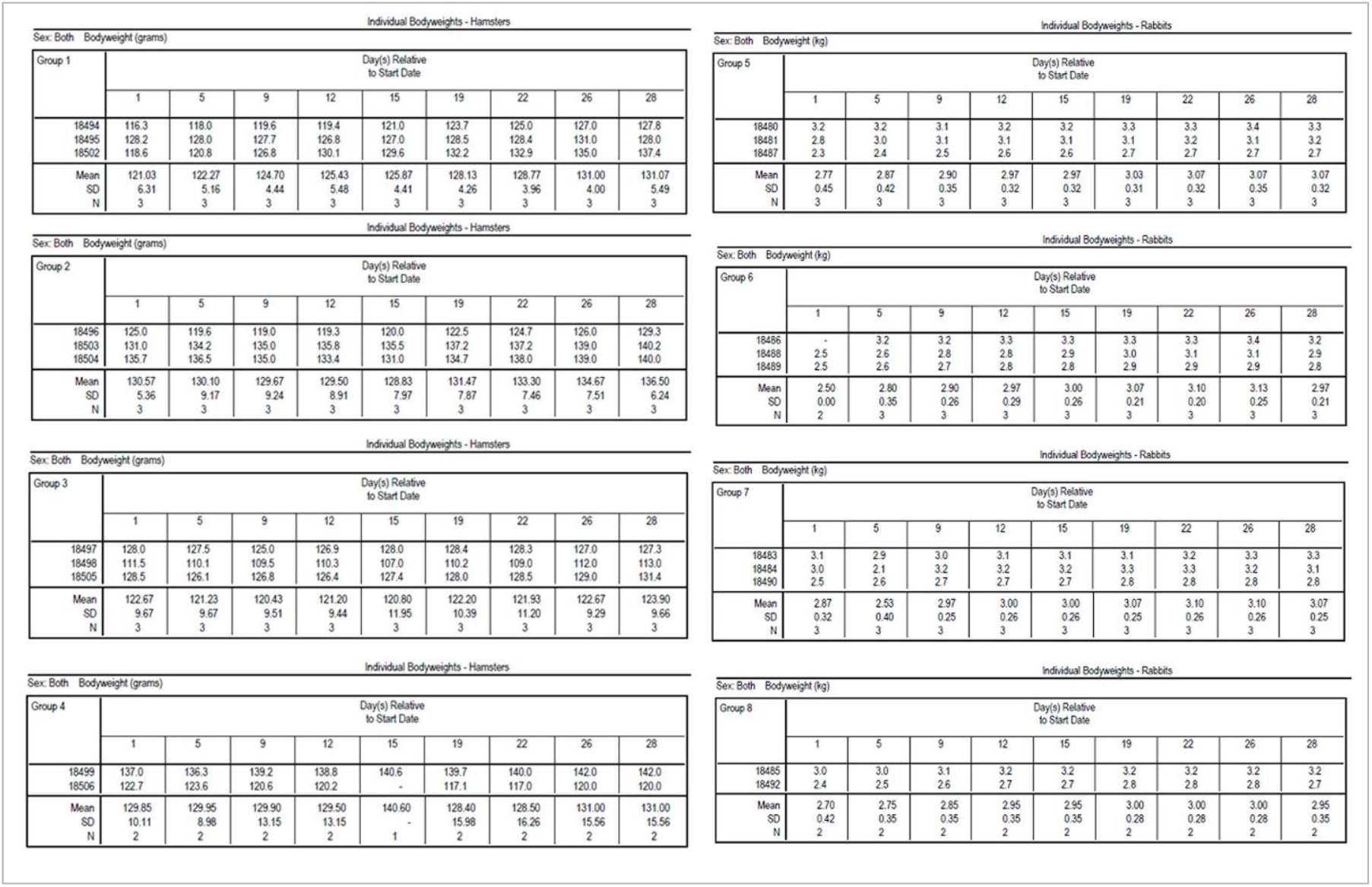
The body weight of rabbits and hamsters monitored throughout the study.

**Table S2.** Draize score and poxvirus gene copy number (*) in rabbits and hamsters monitored during different time points.

| Species |  |  | Day 1 (2 hr post) |  | Day 1 (5 hr post) |  | Day 2 |  | Day 3 |  | Day 4 |  | Day 5 |  | Day 10 |  | Day 20 |  | Day 28 * |
| --- | --- | --- | --- | --- | --- | --- | --- | --- | --- | --- | --- | --- | --- | --- | --- | --- | --- | --- | --- |
|  | Group | Animal | Erythema | Edema | Erythema | Edema | Erythema | Edema | Erythema | Edema | Erythema | Edema | Erythema | Edema | Erythema | Edema | Erythema | Edema | Mean Copy number |
| Hamsters | 1 | 18494 | 2 | 0 | 2 | 0 | 2 | 0 | 2 | 0 | 0 | 1 | 1 | 1 | 1 | 0 | 0 | 0 | 0 |
|  | 1 | 18495 | 1 | 0 | 1 | 0 | 1 | 0 | 1 | 0 | 1 | 0 | 1 | 0 | 1 | 0 | 0 | 0 | 0 |
|  | 1 | 18502 | 1 | 0 | 1 | 0 | 1 | 0 | 1 | 0 | 1 | 0 | 1 | 1 | 1 | 0 | 0 | 0 | 0 |
|  | 2 | 18503 | 1 | 0 | 1 | 0 | 1 | 0 | 1 | 0 | 1 | 0 | 1 | 0 | 0 | 0 | 0 | 0 | 0 |
|  | 2 | 18504 | 1 | 0 | 0 | 0 | 1 | 0 | 1 | 0 | 0 | 0 | 1 | 0 | 0 | 0 | 0 | 0 | 0 |
|  | 2 | 18496 | 2 | 0 | 2 | 0 | 2 | 0 | 1 | 0 | 1 | 0 | 2 | 0 | 1 | 0 | 0 | 0 | 0 |
|  | 3 | 18497 | 1 | 0 | 0 | 0 | 0 | 1 | 1 | 0 | 1 | 0 | 1 | 0 | 1 | 0 | 0 | 0 | 0 |
|  | 3 | 18498 | 1 | 0 | 0 | 0 | 0 | 0 | 0 | 0 | 1 | 0 | 0 | 1 | 0 | 0 | 0 | 0 | 0 |
|  | 3 | 18505 | 1 | 0 | 0 | 0 | 0 | 0 | 0 | 0 | 0 | 1 | 1 | 0 | 0 | 0 | 0 | 0 | 0 |
|  | 4 | 18499 | 1 | 0 | 1 | 0 | 1 | 0 | 1 | 0 | 1 | 0 | 2 | 0 | 1 | 0 | 0 | 0 | 0 |
|  | 4 | 18506 | 1 | 0 | 0 | 0 | 0 | 1 | 0 | 0 | 0 | 1 | 0 | 0 | 0 | 0 | 0 | 0 | 0 |
| Rabbits | 5 | 18480 | 0 | 1 | 0 | 0 | 0 | 1 | 0 | 0 | 0 | 1 | 2 | 1 | 1 | 0 | 1 | 0 | 0 |
|  | 5 | 18481 | 2 | 0 | 2 | 0 | 1 | 0 | 1 | 0 | 2 | 1 | 2 | 1 | 2 | 1 | 1 | 0 | 0 |
|  | 5 | 18487 | 0 | 0 | 0 | 0 | 0 | 0 | 0 | 1 | 2 | 0 | 2 | 0 | 1 | 1 | 1 | 0 | 0 |
|  | 6 | 18488 | 1 | 0 | 0 | 0 | 0 | 1 | 0 | 1 | 0 | 1 | 0 | 1 | 0 | 1 | 0 | 0 | 0 |
|  | 6 | 18489 | 1 | 0 | 1 | 0 | 0 | 0 | 0 | 0 | 0 | 1 | 0 | 1 | 1 | 1 | 0 | 0 | 0 |
|  | 6 | 18486 | 2 | 0 | 2 | 0 | 1 | 0 | 1 | 0 | 2 | 1 | 2 | 2 | 2 | 0 | 1 | 0 | 0 |
|  | 7 | 18483 | 2 | 0 | 2 | 0 | 2 | 0 | 1 | 0 | 1 | 1 | 2 | 0 | 2 | 1 | 1 | 0 | 0 |
|  | 7 | 18484 | 1 | 0 | 1 | 1 | 1 | 1 | 0 | 0 | 0 | 1 | 2 | 0 | 2 | 0 | 1 | 0 | 0 |
|  | 7 | 18490 | 1 | 1 | 1 | 1 | 1 | 1 | 1 | 1 | 0 | 1 | 1 | 0 | 0 | 1 | 2 | 0 | 0 |
|  | 8 | 18485 | 1 | 0 | 1 | 0 | 1 | 1 | 1 | 1 | 2 | 2 | 3 | 1 | 3 | 1 | 1 | 0 | 0 |
|  | 8 | 18492 | 1 | 1 | 1 | 1 | 0 | 1 | 0 | 1 | 2 | 0 | 2 | 1 | 3 | 0 | 1 | 0 | 0 |

## References

1. Kumar, A., et al., Wuhan to World: The COVID-19 Pandemic. Front Cell Infect Microbiol, 2021. 11: p. 596201.

2. Li, J., et al., Data Mining and Content Analysis of the Chinese Social Media Platform Weibo During the Early COVID-19 Outbreak: Retrospective Observational Infoveillance Study. JMIR Public Health Surveill, 2020. 6(2): p. e18700.

3. World Health Organization.14.9 million excess deaths associated with the COVID-19 pandemic in 2020 and 2021. 2022; Available from:https://www.who.int/news/item/05-05-2022-14.9-million-excess-deaths-were-associated-with-the-covid-19-pandemic-in-2020-and-2021.

4. Havlin, J., et al., Immunogenicity of BNT162b2 mRNA COVID-19 vaccine and SARS-CoV-2 infection in lung transplant recipients. J Heart Lung Transplant, 2021. 40(8): p. 754–758.

5. Xu, J., et al., The Cold-Adapted, Temperature-Sensitive SARS-CoV-2 Strain TS11 Is Attenuated in Syrian Hamsters and a Candidate Attenuated Vaccine. Viruses, 2022. 15(1).

6. Xia, S., et al., Safety and immunogenicity of an inactivated COVID-19 vaccine, BBIBP-CorV, in people younger than 18 years: a randomised, double-blind, controlled, phase 1/2 trial. Lancet Infect Dis, 2022. 22(2): p. 196–208.

7. Stephenson, K.E., et al., Immunogenicity of the Ad26.COV2.S Vaccine for COVID-19. JAMA, 2021. 325(15): p. 1535–1544.

8. Dunkle, L.M., et al., Efficacy and Safety of NVX-CoV2373 in Adults in the United States and Mexico. N Engl J Med, 2022. 386(6): p. 531–543.

9. Wouters, O.J., et al., Challenges in ensuring global access to COVID-19 vaccines: production, affordability, allocation, and deployment. Lancet, 2021. 397(10278): p. 1023–1034.

10. Noyce, R.S., et al., Single Dose of Recombinant Chimeric Horsepox Virus (TNX-801) Vaccination Protects Macaques from Lethal Monkeypox Challenge. Viruses, 2023. 15(2).

11. National Research Council (US) Committee for the Update of the Guide for the Care and Use of Laboratory Animals. Guide for the Care and Use of Laboratory Animals; National Academies Press. Eighth Edition ed. 2011: National Academies Press: Washington, DC, USA.

12. Fraser, R., et al., Upper respiratory tract mucosal immunity for SARS-CoV-2 vaccines. Trends Mol Med, 2023. 29(4): p. 255–267.

13. Sunagar, R., A. Singh, and S. Kumar, SARS-CoV-2: Immunity, Challenges with Current Vaccines, and a Novel Perspective on Mucosal Vaccines. Vaccines (Basel), 2023. 11(4).

14. Sheikh-Mohamed, S., et al., Guardians of the oral and nasopharyngeal galaxy: IgA and protection against SARS-CoV-2 infection. Immunol Rev, 2022. 309(1): p. 75–85.

